# Machine learning driven acceleration of biopharmaceutical formulation development using Excipient Prediction Software (ExPreSo)

**DOI:** 10.1101/2025.02.12.637685

**Authors:** Estefania Vidal-Henriquez, Thomas Holder, Nicholas Franciss Lee, Cornelius Pompe, Mark George Teese

## Abstract

Formulation development of protein biopharmaceuticals has become increasingly challenging due to new modalities and higher target drug substance concentrations. The limited amount of drug substance available during development, coupled with extensive analytical requirements, restrict the number of excipients that can be empirically screened. There is a strong need for in silico tools to optimize excipient pre-selection before wet lab experiments. Here, we introduce Excipient Prediction Software (ExPreSo), a supervised machine learning algorithm that suggests excipients based on the properties of the protein drug substance and target product profile. ExPreSo was trained on a dataset comprising 335 regulatory-approved peptide and protein drug products. Predictive features included protein structural properties, protein language model embeddings, and drug product characteristics. ExPreSo showed good performance for the nine most prevalent excipients in biopharmaceutical formulations and minimal overfitting. A fast variant of ExPreSo using only sequence-based input features showed similar prediction power to slower models that relied on molecular modeling. Notably, an ExPreSo variant using only protein-based input features also showed good performance, indicating resilience to the influence of platform formulations. To our knowledge, this is the first machine learning algorithm to suggest biopharmaceutical excipients based on the dataset of regulatory-approved drug products. Overall, ExPreSo shows great potential to reduce the time, costs, and risks associated with excipient screening during formulation development.

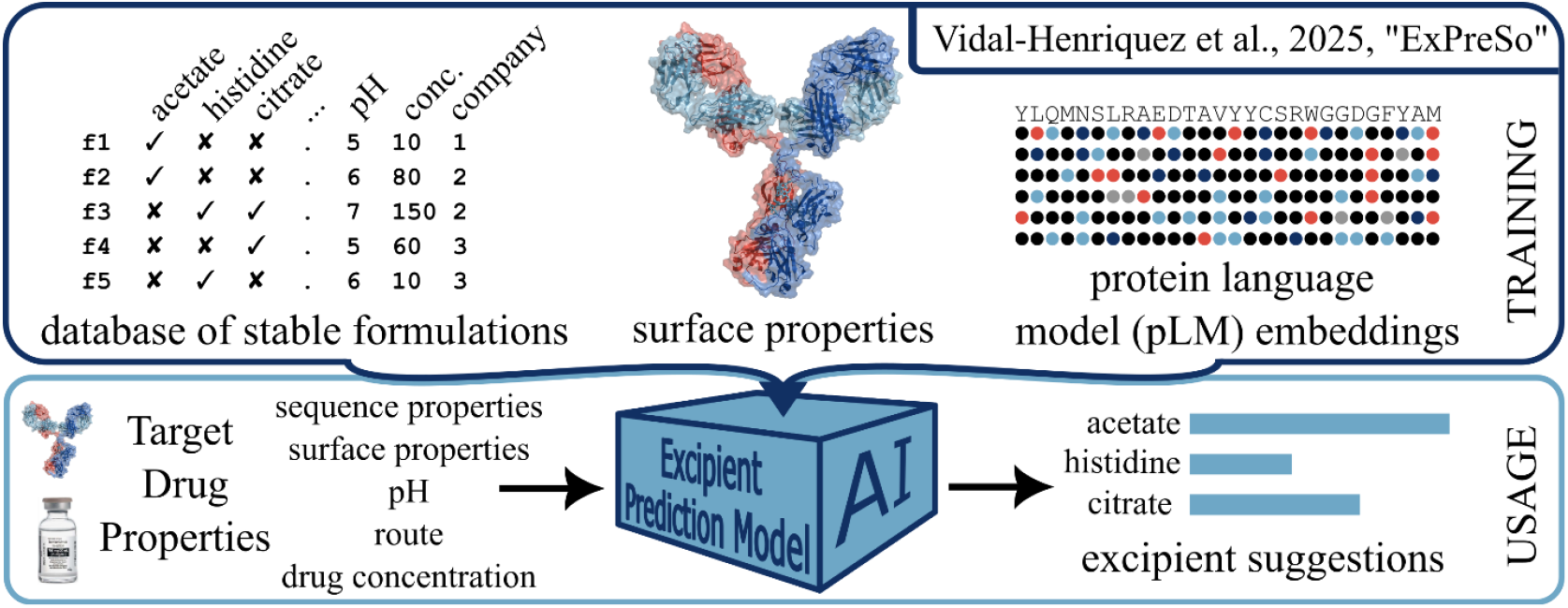

## Introduction

Formulation development is the process in which inactive ingredients are chosen to be added to a drug substance in order to stabilize it for manufacturing, distribution, storage, and patient usage. For protein and peptide biopharmaceuticals, these inactive ingredients, known as excipients, must prevent protein degradation and configuration changes which might affect efficacy and safety.^1^ From a regulatory and operational point of view, it is desirable that formulations have an assigned shelf-life at targeted storage temperature (e.g. 5°C ± 3°C or 25°C ± 2°C at 60% ± 5% relative humidity) of at least 12 months. This has to be enabled by the careful selection of the formulation pH and a limited number of excipients, typically no more than five. The large number of possible excipients and potential instabilities make formulation development a highly complex field. Biologics license applications also need to outline qualitative and quantitative aspects regarding the use of each excipient contained in the drug product (EMEA/CHMP/QWP/396951/2006).

To simplify the challenges of formulation development, some companies utilize *platform formulations* as a starting point for their biopharmaceutical products.^2,3^ Typically, this comprises a standard buffer and a base list of excipients that are tested against individual drug substances. This process is combined with pre-screening to select molecules with a low amount of chemical liabilities and regions prone to aggregation, and presumably, compatibility with the preferred base formulation. Incremental improvements are usually made by exchanging individual excipients with alternatives that have a similar mechanism of action, while keeping the remaining excipients constant. Although this approach saves time and effort^2^, it limits the exploration of a broader range of excipients that might be better suited for the protein of interest.

There is currently a strong interest in the development of biologics drugs with high drug concentrations.^4–7^ This is driven by the increase in biologics with subcutaneous application,^8^ which offers many advantages to patients but usually requires smaller injection volumes when compared to intravenous application. Formulation development of high concentration protein drugs is extremely challenging due to problems with aggregation and viscosity/syringeability.^5^ Furthermore, there are other market trends that offer challenges in formulation development, such as the increasing development of bispecific antibodies^9,10^ and antibody drug conjugates (ADCs).^11^ As a result, more than ever, formulation design must be tailored to the specific characteristics and liabilities of the drug substance under study. This applies to new drug substances under investigation and to the reformulation of existing products, for instance, transitioning from lyophilized to liquid formulations or changing the route of administration from intravenous to subcutaneous.

In the development of biopharmaceuticals, there are *in silico* prediction tools supporting many processes before and after formulation development, but few for the stage of formulation development itself. In early drug development of therapeutic proteins, there are a growing number of *in silico* prediction tools supporting lead development, particularly for antibodies^12^, and for developability assessments aiming to select stable candidates.^13–15^ In later stages of biopharmaceutical development, physics-based methods such as digital twins are well established for real-time process optimization.^16^ However, within the domain of biopharmaceutical formulation development there is a lack of robust, validated *in silico* methods for excipient pre-screening. Molecular docking has been explored to identify excipients binding to a protein,^17^ however most docking algorithms are designed to identify strong binders to act as inhibitors, rather than transient interactions.

The long-term stabilization of a drug substance involves complex stochastic interactions of many elements, such as protein-protein interactions, excipient-protein interactions, and excipient-excipient interactions among others. In order to predict stabilizing excipients using physics-based methods, all these elements need to be modeled simultaneously, for extremely long time scales. Taking all these interactions into account is a computationally intensive task. The computational costs are further increased by the large simulation sizes necessary: for example, most detergents in liquid formulations exist in solution as large micelles, and the simulation size should also be large enough to account for indirect excipient effects such as preferential exclusion (for example by sucrose).^18,19^ So far, this type of calculations have only been published in an academic context for a handful of excipients.^20–24^ Approaches to improve performance include coarse-grained molecular dynamics (MD) simulations,^25^ AI powered MDs that utilize machine learning-refined force-fields,^26^ and machine learning models trained to predict the outcome of MD simulations.^27^

An alternative strategy to predict excipient binding *in silico* is to conduct short all-atom MD simulations with the protein surrounded by a high concentration of the desired excipient. This technique, known as fragment mapping, can be used to rank excipients according to their affinity to a protein.^28–31^ Every method that looks at overall protein-excipient interactions relies on the assumption that excipients with increased protein binding are also more likely to increase protein stability. However, it has been shown that in many cases stronger excipient binding does not improve stability.^32^ A more targeted strategy would involve screening excipients predicted to bind the exact sites of protein-protein interaction or transient unfolding that lead to aggregation, but these critical regions are usually unknown.

An alternative to physics-based modeling is a knowledge-based approach, particularly the development of machine learning algorithms to predict stable formulations. The primary challenge with such methods lies with data availability, as effective machine learning models require diverse datasets encompassing a large number of excipients and drug substances. While several machine learning algorithms have been created to assist small molecule formulations, such as those involving solid dispersion and cyclodextrin formation^33,34^, no such equivalent exists for biologics. Since formulation data during drug development is kept confidential, published datasets are very small and few companies have a sufficiently large and diverse drug substance portfolio to support machine learning development. On the other hand, the final formulations of biopharmaceutical drugs approved by regulatory authorities are publicly accessible. This database of formulations is growing rapidly^7,35^ and it has already enabled the first quantitative analyses of stabilizing excipients,^7,35–37^ and trends over time.^3^ To our knowledge, this data remains an untapped resource for the prediction of stabilizing excipients.

This machine learning approach is further empowered by recent advances in computational tools such as AlphaFold2 and protein language models (pLMs). AlphaFold2 is a machine learning algorithm that provided a breakthrough in the de-novo prediction of protein structures.^38^ These improved structural predictions enable the extraction of more reliable protein properties, which serve as inputs for downstream machine learning applications. pLMs are large language models that have been trained explicitly to predict protein sequences. A byproduct of pLMs are protein embeddings, which are the vectorial representations in the language model of each amino acid in a sequence.^39^ Once a pLM is trained, the protein embeddings can be rapidly generated from any input protein sequence. These embeddings encode information about the amino acid and their surroundings in the sequence. Their use has provided a leap forward in predictive power for different machine learning tasks such as the prediction of structure, function, and epitopes.^40–43^

In this study, we created the Excipient Prediction Software (ExPreSo), a set of machine learning models trained on a database of formulations of 335 approved biopharmaceutical products. Each model predicts the probability that a specific excipient is included in a stable formulation, given inputs such as the drug substance sequence, pH, stock keeping unit (liquid or lyophilized), and drug substance concentration. ExPreSo has predictive power for nine commonly used excipients, offers interpretability regarding the most important predictive features, and can generate results within seconds, enabling its use in early-stage formulation development.

## Materials and Methods

A comprehensive database was made of all formulations of FDA-approved drug products as of 27 September 2024. This database was filtered to include formulations with only one active pharmaceutical ingredient for which an amino acid sequence was available. The formulations were then converted to a binary matrix, with rows corresponding to individual formulations, and columns to excipient presence. The excipient names were harmonized by unifying compounds that are identical in solution, such as mono- or di-basic forms of a buffer, or their hydrated and anhydrous variants. For example, formulations were deemed to contain the excipient ‘sodium phosphate’ whether the listed ingredient was the mono- or dibasic form, or whether the powdered form of the chemical compound was hydrated or anhydrous. The matrix entries were Boolean (True or False), denoting the presence or absence of each excipient in a formulation. Duplicate formulations were removed if they had the same International Nonproprietary Name (core name, excluding biosimilar suffixes) and excipients. Excipients found in less than 10% of the formulations were then removed, resulting in nine remaining excipients. A final round of duplication removal was performed to eliminate any redundancies caused by the reduced excipient set. This process yielded a dataset of 335 unique biopharmaceutical formulations.

To this dataset of formulations, we added information on the drug product. Categorical variables, including the route of administration and stock keeping unit (liquid/lyophilized) were added using one-hot encoding. To manage the diversity of marketing authorization holders, the most commonly appearing companies were retained as individual categories, while the remaining companies were grouped under a collective “other company” label. The number of kept companies was selected such that the “other company” category amounted to approximately 50% of the total formulations. Numerical features including the product’s marketing start date, the formulation pH, and the drug concentration were retained in their original form. Any missing numerical value was imputed using the mean of the available data.

We then augmented the dataset with a range of protein based features (descriptors) derived from the primary amino acid sequence. For each protein, we calculated the frequency of each individual amino acid and each possible dipeptide pair in the sequence. Sequence-based physicochemical properties, including isoelectric point and fraction of helix, sheet, and turn residues, were calculated using the BioPython ProtParam module.^44^ Advanced protein representations were also generated via protein language model embeddings using the ProtTrans prot_t5_xl_half_uniref50 model^42^ through the provided prott5_embedder.py script. For proteins with multiple polypeptide chains, sequences were concatenated prior to all calculations. ProtTrans pLM embeddings are a 1024-dimensional vector for each amino-acid in the protein sequence, which we aggregated for each protein by computing the mean vector along the entire sequence, resulting in one 1024-dimensional vector per sequence.

The dataset was further enriched with features derived from generated 3D protein structures. Monoclonal antibody (mAb) structures were generated using the antibody modeler algorithm of Chemical Computing Group Inc Molecular Operating Environment (MOE) version 2022.02, with the default Fc glycosylation. Non-mAb drug substances were modeled using AlphaFold2^38^ and subsequently protonated using MOE’s QuickPrep protocol. Surface properties at pH 7.0 and 0.1 M NaCl were calculated with MOE. Drug substances were classified as mAbs if a complementarity-determining region (CDR) was detected by MOE, and the IgG heavy chain subtype (IgG1, IgG2 or IgG4) was represented using one-hot encoding. The full list of input features is given in Supplementary Table S1. While the code and full dataset are proprietary, a reduced dataset with formulations marketed prior to 2020, including all machine learning features, is included in the Supplementary Data.

The ExPreSo pipeline, including the nested validation procedure, is presented as a pseudoalgorithm in the Supplementary Data. Four distinct models were created, each with a different subset of input features: All Features, Interpretable, Fast, and Protein-Based. The Interpretable model excluded amino acid and dipeptide frequency and pLM embedding features. The Fast model omitted molecular modeling-derived features to enable rapid computation. The Protein-Based model contained only features intrinsic to the protein, such as protein type, molecular modeling-derived surface properties, dipeptide frequencies, and pLM embeddings; and lacked information such as the pH, company, drug substance concentration, and whether or not the product was lyophilized.

Given that the initial number of features (1582 in the All Features model) was far higher than the number of formulations (335), feature reduction was applied to each model separately to prevent overfitting. We first removed highly correlated features (absolute Pearson R>0.8), reducing the feature counts to 865, 81, 827, and 831 for the All Features, Interpretable, Fast, and Protein-Based models, respectively. We then applied principal component analysis (PCA) to reduce the number of features to ensure there were less features than independent observations, while retaining most of the variance. To maintain interpretability, PCA was applied separately within manually defined feature groups. We retained ten dimensions (features) for amino acid frequency, 32 for dipeptide frequency, and 32 for pLM embeddings. For the structural features derived from molecular modeling, related properties were first assigned to one of six groups by similarity, and then reduced to two dimensions per group via PCA. These groups are listed in Supplementary Table S2 along with the fraction of covariance explained by the PCA during feature reduction. Features that did not belong to a group were left unchanged. This process reduced the original set of 89 MOE-derived properties to 39, thereby reducing noise while retaining model interpretability. The final feature counts were 141, 67, 119, and 107 for the All Features, Interpretable, Fast, and Protein-Based models respectively. To prevent information leakage, the PCA transformers were trained exclusively on the train set, and applied later to both the train set and blind test set as required.

The nine different machine learning algorithms were trained independently of each other using the ensemble Extra-Trees algorithm^45^ as part of the python Scikit-learn package version 1.5.2 using the default hyperparameters.^46^ The Extra-Trees algorithm is very similar to Random Forest, but introduces additional randomness by selecting split thresholds for each feature completely at random, rather than by optimized splits.^45^ This results in faster training than Random Forest but maintains a robust calculation of feature importances, a capability recently leveraged to investigate therapeutic protein developability.^47^ To address class imbalance in the dataset we performed SMOTE oversampling^48^ using the imblearn library with default settings.

Formulations from highly similar proteins were grouped into clusters to ensure they were never present in both train and test sets during machine learning validation. Sequence-based clustering was performed using CD-HIT^49,50^ on concatenated sequences containing all domains of the full protein, with identity thresholds set at 95% for mAbs, and 80% for non-mAbs. Formulations were also clustered together if the drug substances shared an INN. This strategy ensured that biosimilars were grouped with their reference innovator products, antibody-drug-conjugates with their unconjugated forms, and all peptide variants (e.g. insulin human, insulin aspart, insulin glargine) within the same cluster. The 335 non-redundant formulations were assigned to 192 clusters. During machine learning validation, all members within a cluster were either part of the train set or the test set, but were never split between both.

Machine-learning validation was performed using ten iterations of Monte Carlo cross validation (MCCV) with an inner loop of leave-one-group-out cross validation (LOGO-CV). In each MCCV iteration, a blind test set was generated by randomly selecting 20% of the clusters (without stratification), while the remaining formulations comprised the train set. Each MCCV iteration used a different random seed that affected selection for the test set, SMOTE oversampling, and model training. In each MCCV iteration, the same train/test split was used for all excipients and models (feature sets). To ensure robust evaluation, the blind test set included at least two formulations containing each excipient, and at least two formulations lacking each excipient. To prevent overrepresentation by drugs that have been extensively reformulated, only two randomly chosen representatives from each cluster were retained in the blind test set, and the remainder were dropped. For each excipient target in each MCCV iteration, we retained only the top 20 predictive features according to SHAP.^51^ Afterwards, cross-validation was conducted using an inner loop of LOGO-CV, where the algorithm was trained iteratively on all clusters except one, and then tested on the remaining cluster. The final blind test validation for each iteration and excipient target was performed using a model trained on the full train set. Unless otherwise specified, all performance metrics (e.g. ROC-AUC, PR-AUC, overfitting index) represent the mean of ten MCCV iterations.

## Results and Discussion

We curated a comprehensive dataset of 335 FDA-approved formulations of protein and peptide biopharmaceuticals, representing 241 unique drug substances. Monoclonal antibodies (mAbs) constituted 45% of the dataset, with IgG1 subtypes being the most prevalent (30% of the dataset, 68% of the mAbs, Figure 1A). An analysis of the FDA approval dates shows that the number of protein/peptide drug formulations is increasing rapidly (Figure 1D), with a large proportion of our dataset being approved in the last 5 years. The dataset included relevant information on formulation attributes, including excipients, whether it was liquid or lyophilized, the year of approval, and the company (marketing authorization holder). To support machine learning, we supplemented the dataset with features derived from the *in silico* modeled 3D structures of the drug substances, such as size of surface patches (positive, negative, ionic, hydrophobic), charge dipole, and isoelectric point (Table S1). The dataset was also supplemented with sequence-based features, including amino acid frequency, dipeptide frequency, and protein language model (pLM) embeddings extracted from the ProtT5 model.^42^.

**Figure 1.**
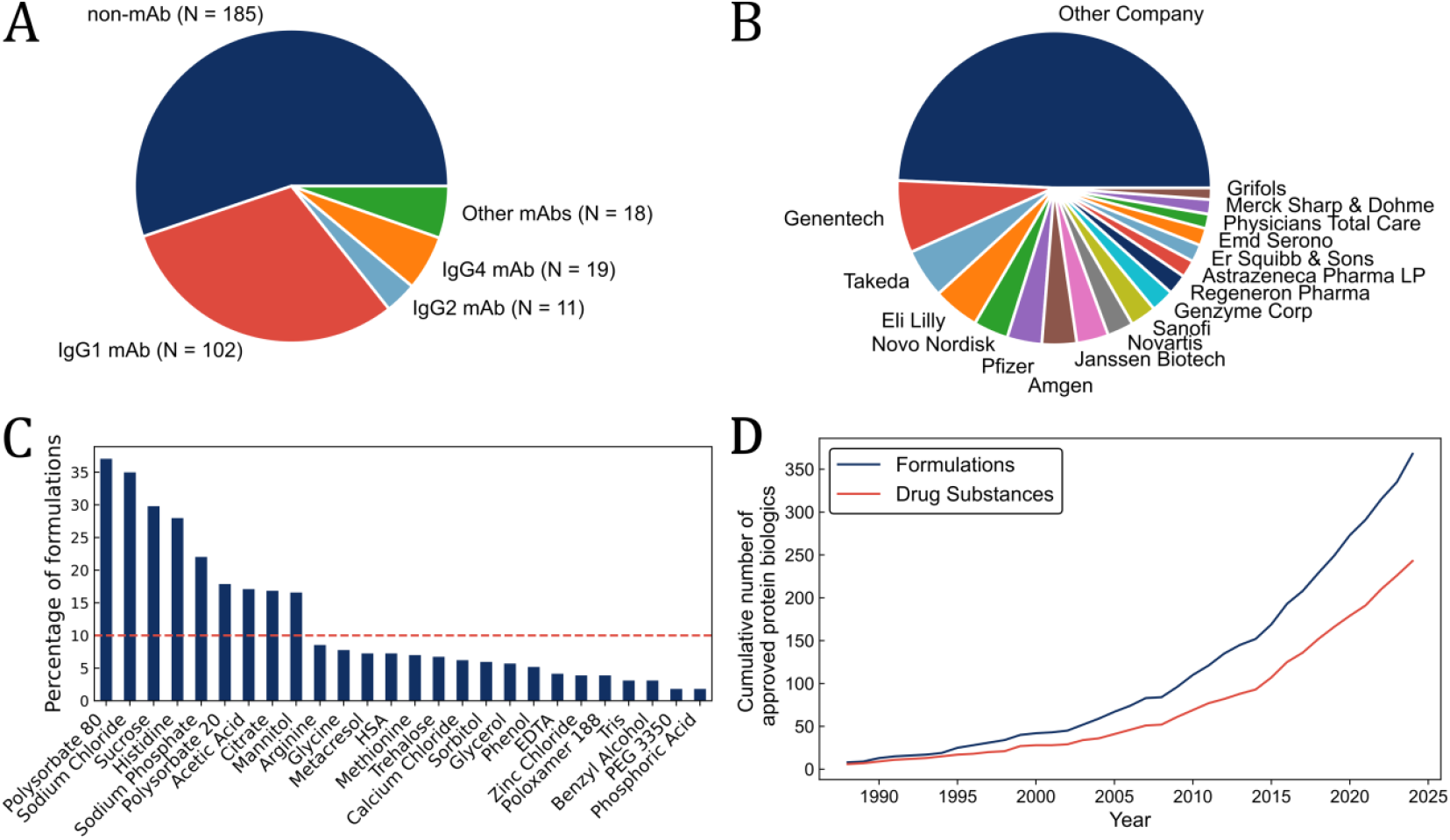
Main characteristics of the dataset used for machine learning. A) Distribution of antibody heavy chain subtypes. B) Companies retained in the input features. The large “Other Company” category enables the use of this feature in ExPreSo as an independent bioinformatic tool. C) Abundance of excipients in the dataset prior to selecting the top excipients used in ExPreSo and removal of duplicate formulations (see Methods). D) Cumulative number of approved formulations and drug substances in the ExPreSo dataset, before duplicate removal (same data as in Figure 1C).

To develop an interpretable predictive tool for excipients in formulations, we implemented a collection of independent machine learning algorithms, each focused on a single excipient. Specifically, we employed the ExtraTrees classifier^45^, an algorithm similar to Random Forest^52^, to predict the presence or absence of an excipient in stable final drug products. All excipient prediction models used the same set of predictive features, which included protein properties and target product profile. To allow de-novo formulation suggestions, the predictive features did not contain any information regarding the presence of other excipients in the stable formulation. Preliminary experiments revealed that prediction power was only reliable for excipients that occurred frequently in the dataset, therefore ExPreSo is limited to excipients present in more than 10% of formulations. This yielded nine excipient targets: acetic acid, citrate, histidine, mannitol, polysorbate 20, polysorbate 80, sodium chloride, sodium phosphate, and sucrose.

ExPreSo had the highest prediction power for acetic acid, histidine, and polysorbate 80, and prediction for all excipients was better than a random predictor (Figure 2). With the exception of acetic acid, there was a correlation between predictive power and the abundance of the excipient in the dataset (R^2^ = 0.54 without acetic acid, 0.13 with acetic acid, see Figure 2C). This was not unexpected, as the more common excipients (polysorbate 80, sodium chloride, sucrose, and histidine) had a more balanced dataset and thus gave models with better prediction power. Conversely, less common excipients (citrate, mannitol, and polysorbate 20) tended to exhibit lower prediction metrics.

**Figure 2.**
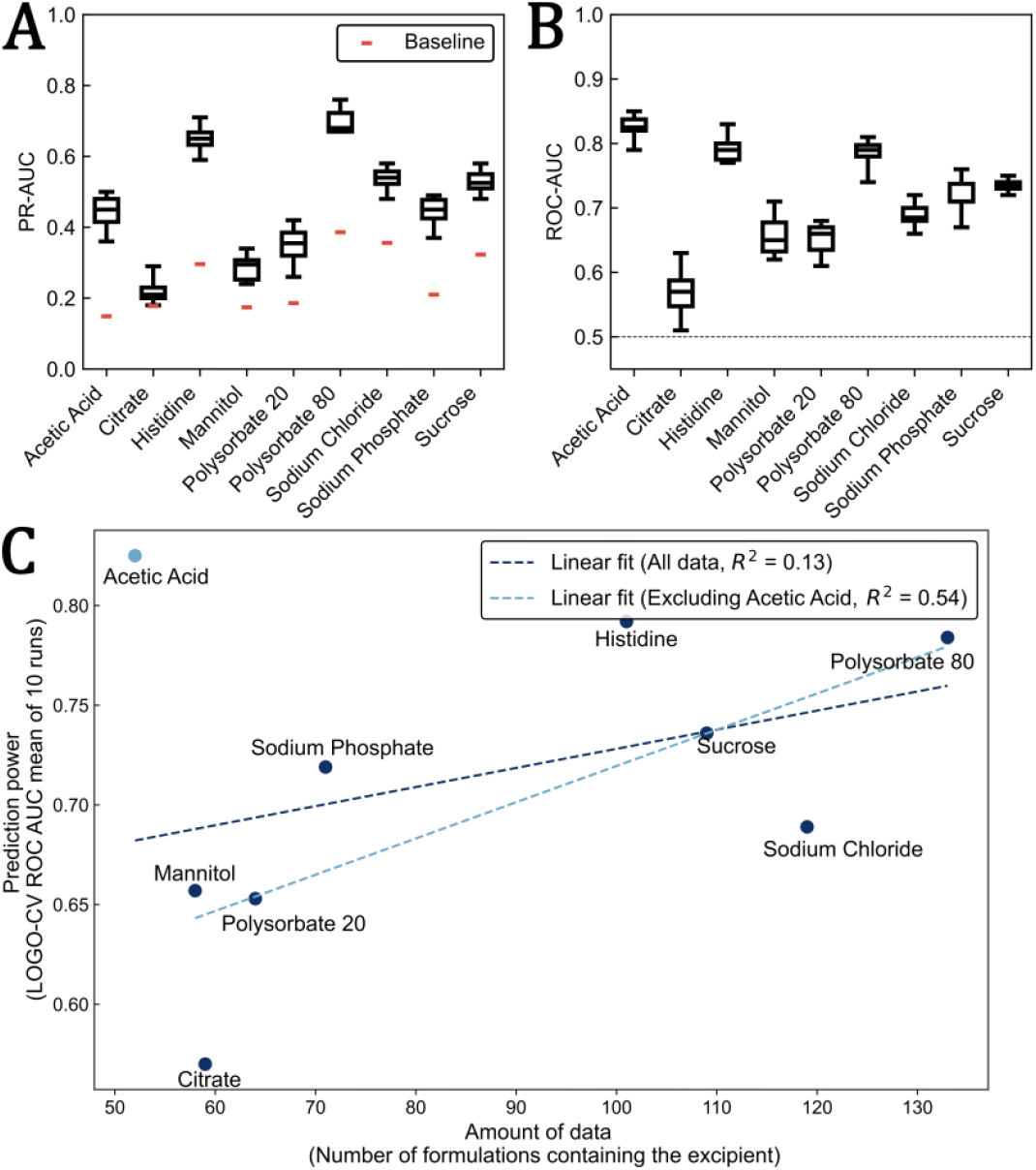
Predictive performance of ExPreSo for nine excipients. Model performance was evaluated using leave-one-group-out cross-validation on the train set with all predictive features (“All Features”). Results were aggregated over 10 Monte Carlo cross-validation (MCCV) runs, each using a different random seed for test set creation. In each boxplot, whiskers indicate min and max, and the central line indicates the median. A) Precision Recall Area Under the Curve (PR-AUC). The baseline (red dash) indicates the precision of a random predictor. B) Receiver Operating Characteristic Area Under the Curve (ROC-AUC). The baseline (dotted line) indicates the performance of a random predictor. C) ROC-AUC prediction power showed correlation to excipient abundance, with the exception of the acetic acid outlier. Each point represents the mean ROC-AUC from 10 MCCV runs.

We validated the predictive power of ExPreSo using a leave-one-group-out cross validation (LOGO-CV) methodology and a blind test validation (see Methods). Model performance was quantified using area under the curve (AUC) for the Receiver Operating Characteristic (ROC) and for the Precision-Recall (PR) curves. The dataset was small and unbalanced, with substantially more formulations lacking each target excipient than containing it. Because of this, there was high run-to-run variation in prediction performance, particularly for the blind test dataset. To counteract this we implemented a Monte Carlo cross validation (MCCV) strategy, averaging performance metrics for 10 runs with different random seeds. Using this approach, ExPreSo consistently showed high prediction power (ROC-AUC > 0.7) for 5/9 excipients in the LOGO cross-validation.

While some excipients performed consistently over several runs, such as polysorbate 80 and histidine (see Figure 2), other excipients such as citrate and mannitol showed a high variation in model performance. We attribute this inconsistency to the limited size of the dataset and how formulations are allocated to the train and test set in each run. This underscores the challenge in achieving stable predictions for less represented excipients.

The ExPreSo algorithm is adaptable and can be configured for either interpretability or computational speed, depending on the chosen subset of input features. For the “Interpretable” model, we excluded sequence-based features such as pLM embeddings, amino acid frequency, and dipeptide frequency. The retained structure-derived protein features are more readily human interpretable when examining feature importances. To enable ExPreSo as a standalone predictor, we created a “Fast” model that excludes features derived from the computationally expensive molecular modeling, and instead relies on rapidly computable sequence-based features such as pLM embeddings and dipeptide frequencies. While the Fast model allows predictions to be calculated in milliseconds, the feature importances have low human interpretability because pLM embeddings encapsulate complex, abstract representations, and dipeptide frequencies have limited direct biological relevance. The performance of the All Features, Interpretable and Fast models were comparable (Figure 3).

**Figure 3.**
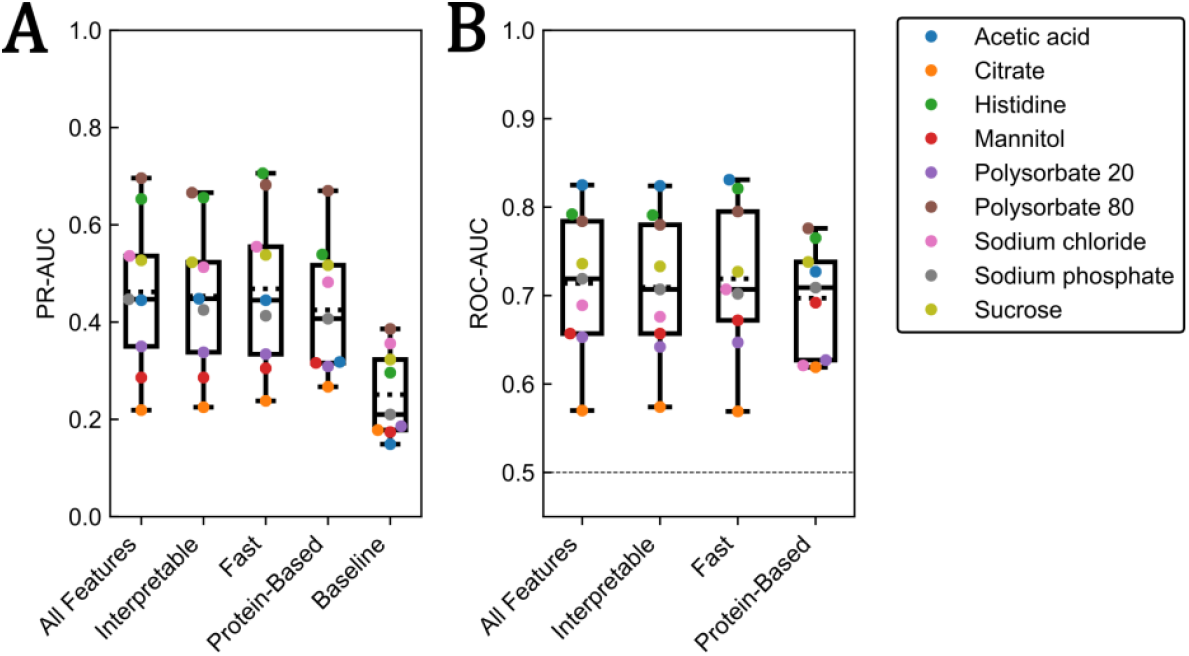
All Features, Fast, Interpretable, and Protein-Only versions of ExPreSo show similar performance. ExPreSo was validated using four models, each with a different subset of input features (descriptors). The “Interpretable” model lacked sequence-based features such as pLM embeddings. The “Fast” model lacked features derived from molecular modeling. The “Protein-Based” model used only features based on the protein sequence and 3D structure. Shown are the area under the curve (AUC) metrics from leave-one-group-out cross-validation (LOGO-CV), averaged over ten Monte Carlo cross-validation (MCCV) runs (see Methods). Boxplots present Area Under the Precision Recall Curve (PR-AUC) and Area Under the Receiver Operator Characteristic Curve (ROC-AUC); the solid middle line indicates the median, whereas the dotted line indicates the mean. For the ROC-AUC, the baseline (thinner dotted line at 0.5) indicates the performance of a random predictor.

To better understand the potential bias of platform formulations, where companies use the same base formulation for multiple products, we analyzed the similarity of formulations within each company using the Jaccard similarity index. The Jaccard index compares how similar two sets are. It is defined as the number of elements in the sets’ intersection divided by the number of elements in their union. This metric allowed us to systematically assess formulation redundancy. Surprisingly, our analysis revealed that only a small fraction of the formulation pairs exhibited high similarity, both when considering the entire dataset (Supplementary Figure S2), and the top companies separately (Supplementary Figure S3). These finding indicate that although many companies may start with a base formulation, they tend to refine each final formulation to the specific needs of the given drug substance.

To further assess whether company-specific practices were influencing ExPreSo predictions, we constructed a “Protein-Based” model that excluded all features related to the manufactured product, such as the company, pH, year of approval, and route of administration. Instead, the model incorporated only features associated with the protein drug substance, including surface properties derived from molecular modeling, protein type, pLM embeddings, and dipeptide frequencies. If there were a strong platform formulation bias, we would anticipate a substantial drop in predictive performance when compared to models with the full feature set. Interestingly, the Protein-Based model showed only a small decrease in predictive power (Figure 3), showing that the dataset and algorithm are surprisingly resilient to the bias of platform formulations. Together with the low Jaccard index values of formulation similarity, this supports the conclusion that excipient selection is generally more tailored to the drug substance than previously assumed. Therefore, there are rational reasons for a company’s excipient bias, such as the stabilizing effect of these excipients for a favored protein scaffold, or the selection of drug candidates that are stable in the preferred platform formulation, as part of a ‘platform fit’ approach.^3^

Overfitting is a common issue in machine learning models trained on small datasets with many input features. To minimize overfitting, we implemented a number of measures during the development of ExPreSo. Overfitting typically manifests as much higher metrics in cross-validation (where some data are seen in both training and testing) compared to a true hold-out “blind” test set, whose observations are entirely unseen during training. For models with a single prediction target, overfitting is often assessed qualitatively by plotting validation metrics, such as the ROC, for the cross-validation and blind test sets. However, because ExPreSo evaluates nine different prediction targets (excipients) across four different model types (All Features, Interpretable, Fast, Protein-Based), uses two different validation metrics (ROC-AUC, PR-AUC), and averages performance over 10 MCCV runs to reduce variability, traditional visualization approaches proved impractical. We therefore developed an “overfitting index”, defined as the difference between the cross-validation and blind test performance metrics, normalized to the value of the cross-validation metric:

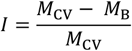

where *I* is the overfitting index, *M*_CV_ is the performance metric from cross-validation, and *M*_B_ is the performance metric from a blind test set. We use *M*_CV_ as a baseline because it is much more consistent than *M*_B_ for small datasets. A higher overfitting index is undesirable as it indicates a greater discrepancy between metrics for training and truly unseen data, indicating poor predictive power towards new observations. For example, a ROC-AUC of 0.7 after cross-validation and 0.6 for the blind test set yields an overfitting index of (0.7-0.6)/0.7 ≈ 0.14, which can be loosely interpreted as a 14% overestimation of model performance in cross-validation.

We applied several techniques to reduce overfitting, including the automatic removal of correlated features, feature reduction using principal component analysis (PCA), and limiting all models to use only the 20 most predictive features (see Methods). After these refinements, all algorithm variants demonstrated minimal overfitting (Figure 4). Specifically, the difference in performance between LOGO-CV and blind test sets was less than 25% for all excipients, with a median difference below 10%. This narrow gap between validation and blind test metrics supports the conclusion that the model maintains strong predictive power when applied to new drug substances.

**Figure 4.**
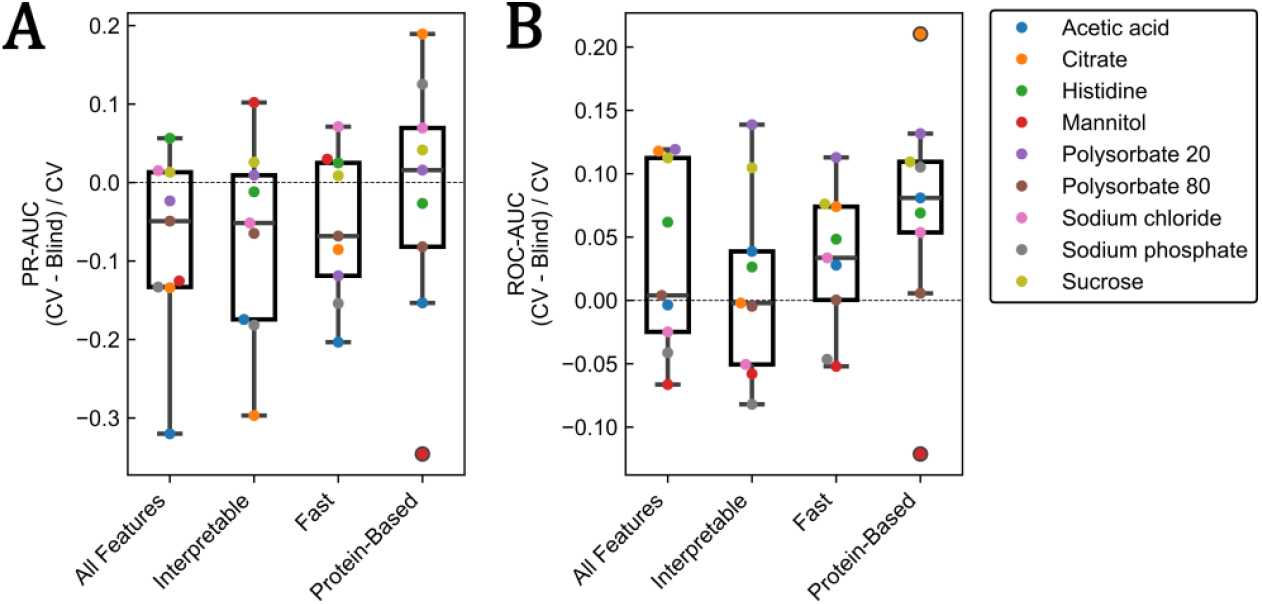
Minimal overfitting was observed across all model variants. “Overfitting index”, defined as the relative difference in performance between the LOGO-CV and the blind test set. Higher values indicate larger overfitting. Values above zero (dotted line) indicate better performance for the LOGO-CV than the blind test. The central line in each boxplot corresponds to the median; box edges indicate the interquartile range, and whiskers extend to datapoints within 1.5 times the interquartile range. Results represent the mean values across ten repeated Monte Carlo cross-validation (MCCV) runs, as described in the Methods.

Examination of feature importances in the Interpretable model revealed that both the target product profile and protein features were important for prediction (see Supplementary Figure S1). Many of the most influential features are proxies for protein types, such as length, mass, and percentage of beta sheet residues, suggesting that the model was appropriately distinguishing mAbs from non-mAbs. As the dataset grows and when a more stringent clustering of similar proteins can be performed we expect that feature importances would indicate more directly the surface properties that are relevant to excipient selection.

Interestingly, our analysis indicated that acetic acid showed the highest predictability among the excipients, despite having the least data available (Figure 2C). We propose that the presence of acetic acid in approved formulations is highly predictable due to its complete absence in lyophilized formulations, and association with low pH. This hypothesis is further supported by the marked reduction in prediction performance for acetic acid in the Protein-Based model, which lacked both the formulation type and pH as input features.

Algorithms such as ExPreSo can be used to provide excipient suggestions, in order to guide screening experiments during formulation development. It is critical, however, that the prediction scores from ExPreSo are interpreted correctly. Specifically, ExPreSo predicts the likelihood that a given drug substance would be formulated with a particular excipient, if it were in a drug product within this historical dataset and with the input target product profile. Consequently, ExPreSo’s outputs could be included as one factor among many in excipient preselection, alongside others such as pre-formulation experimental findings, stable formulations for similar molecules, and the latest scientific literature relevant to the modality, protein family, and individual excipients under consideration.

A possible improvement for ExPreSo would be to instead of having a simple classifier, to create a multi-class predictor. Which would instead of independently assessing the presence or absence of a single excipient (e.g. polysorbate 80), would predict the assignment to one of several exclusive classes (e.g. polysorbate 80, polysorbate 20, poloxamer 188, or no detergent). However, implementing such a system poses significant challenges: biopharmaceutical formulations frequently include multiple excipients from the same class (particularly buffers or sugars/polyols), and individual excipients often perform multiple roles (such as histidine acting as both an amino acid and a buffer), complicating class assignment. As the number of marketed antibody-drug conjugates increases, another area for improvement would be the incorporation of features representing the properties of the attached linkers and payloads, rather than restricting the input features to only the protein component. Extending the algorithm to include formulations containing multiple proteins, currently excluded for simplicity, would also broaden the algorithm’s applicability. The most significant improvement, however, would be a substantial expansion of the dataset. Because of the high failure rate of drug candidates during preclinical and clinical development, the dataset of approved drugs represents only a small fraction of stable therapeutic protein formulations in existence. The formulations of drugs undergoing clinical trials are undisclosed. Increasing the availability of the data currently held by individual pharmaceutical companies would improve the efficiency of biopharmaceutical formulation as a whole, reduce the costs and risks associated with drug development, and bring great benefits to patients worldwide.

When interpreting ExPreSo results to design experiments it is important to consider the underlying assumption that the presence of each excipient is independent from one another. As the models lack information on the presence of other excipients, they cannot predict complete, compatible, stabilizing formulations. For example, the top ranked excipients might include excipient pairs that share a similar mode of action (e.g. polysorbate 80/polysorbate 20 or sucrose/trehalose) and should not be combined in a single formulation. When predicting these excipient pairs, ExPreSo might suggest testing either or both of them. Therefore, we recommend to use machine learning-derived suggestions as a starting point for a multi-factor or design of experiments (DoE) screening experiment, and to analyze the logical sense of each excipient suggested by ExPreSo before starting experiments.

ExPreSo-like algorithms could also be modified to support later stage excipient screening scenarios where some excipients (e.g. buffer and detergent) are already fixed. In such cases, incorporating information on selected or excluded excipients as input features would allow the model to simultaneously assess protein-excipient and excipient-excipient compatibility.

Another possible application could be to assess the potential fit of a drug substance candidate to established platform formulations during developability screening. For instance, if the target platform formulation includes histidine, sucrose, and polysorbate 80, the combined ExPreSo predictions for these three excipients could be aggregated to create a metric to assess developability. Selecting candidates that suit an established formulation could bring cost savings in formulation development and manufacturing.^3^

Finally, it is important to highlight that ExPreSo currently predicts only the likelihood of excipient presence, not their optimal concentrations. As a result, it cannot substitute thorough experimental testing of different excipient concentration combinations to identify the most effective formulation.

## Conclusion

To our knowledge, ExPreSo is the first machine learning algorithm to suggest excipients for biopharmaceutical formulation development using a dataset of regulatory-approved drug products. In a simple and interpretable approach, we created a group of independent machine learning classifiers. Each algorithm predicts whether a particular excipient would be included in a stable formulation for the given drug substance. The algorithms had good predictive performance with minimal overfitting. Extensive validation confirmed they can accurately propose excipients for new drug products entering the market. ExPreSo also proved surprisingly resilient to the influence of platform formulations in the dataset, although its predictive power improved slightly when additional product details such as company were provided as input features. The Fast version of ExPreSo can generate predictions within seconds.

ExPreSo assists excipient preselection by enabling objective, data-driven decision making, thereby helping to reduce human bias. In the future, as AI-based excipient suggestion algorithms continue to improve, it should be possible to reduce the number of excipients screened during formulation development. This will ultimately lower costs for pharmaceutical companies and patients, while also shortening drug development timelines. Since ExPreSo’s performance depends on the volume and diversity of approved drug formulation data (Figure 2C), the ongoing increase in new biologics entering the market (Figure 1D) will considerably expand its dataset. This expansion will enhance ExPreSo’s predictive accuracy and broaden the spectrum of excipients it can reliably suggest, boosting its overall utility. Additionally, with a curated dataset containing excipient concentrations, future versions of ExPreSo could be extended to predict excipient concentrations, offering even more comprehensive guidance to formulation scientists.

## Abbreviations used

ExPreSo: Excipient Prediction Software
ROC: Receiver Operating Characteristic
AUC: Area Under the Curve
TPP: Target Product Profile
FDA: Food and Drug Administration
PCA: Principal Component Analysis
LOGO-CV: Leave-One-Group-Out Cross Validation
mAb: monoclonal antibody
CDR: Complementarity-Determining Region
SHAP: SHapley Additive exPlanations
MOE: Molecular Operating Environment

## Acknowledgments

We would like to thank Dmitrij Frishman of Technische Universität München for helpful feedback on the bachelor thesis of Nicholas Lee in the initial stages of the project. We would also like to thank Theodore W. Randolph of University of Colorado Boulder, Andreas Seidl, and Nehil Chaturvedi for helpful comments.

Statement: During the final stages of preparing this work, the authors used ChatGPT-4.1 in order to improve text readability of the manuscript. After using this tool, the authors reviewed and edited the content as needed and take full responsibility for the content of the publication.

## References

1. Akers MJ. Excipient-drug interactions in parenteral formulations. J Pharm Sci. 2002;91(11):2283–2300. doi:10.1002/jps.10154

2. Warne NW. Development of high concentration protein biopharmaceuticals: The use of platform approaches in formulation development. Eur J Pharm Biopharm. 2011;78(2):208–212. doi:10.1016/j.ejpb.2011.03.004

3. Mieczkowski CA. The evolution of commercial antibody formulations. J Pharm Sci. 2023;112(7):1801–1810. doi:10.1016/j.xphs.2023.03.026

4. Desai M, Kundu A, Hageman M, Lou H, Boisvert D. Monoclonal antibody and protein therapeutic formulations for subcutaneous delivery: high-concentration, low-volume vs. low-concentration, high-volume. MAbs. 2023;15(1). doi:10.1080/19420862.2023.2285277

5. Zarzar J, Khan T, Bhagawati M, Weiche B, Sydow-Andersen J, Alavattam S. High concentration formulation developability approaches and considerations. MAbs. 2023;15(1). doi:10.1080/19420862.2023.2211185

6. Jiskoot W, Hawe A, Menzen T, Volkin DB, Crommelin DJA. Ongoing challenges to develop high concentration monoclonal antibody-based formulations for subcutaneous administration: Quo Vadis? J Pharm Sci. 2022;111(4):861–867. doi:10.1016/j.xphs.2021.11.008

7. Ghosh I, Gutka H, Krause ME, Clemens R, Kashi RS. A systematic review of commercial high concentration antibody drug products approved in the US: formulation composition, dosage form design and primary packaging considerations. MAbs. 2023;15(1). doi:10.1080/19420862.2023.2205540

8. Prašnikar M, Bjelošević Žiberna M, Gosenca Matjaž M, Ahlin Grabnar P. Novel strategies in systemic and local administration of therapeutic monoclonal antibodies. Int J Pharm. 2024;667(May):0–2. doi:10.1016/j.ijpharm.2024.124877

9. Holmes D. Buy buy bispecific antibodies. Nat Rev Drug Discov. 2011;10(11):798–800. doi:10.1038/nrd3581

10. Spiess C, Zhai Q, Carter PJ. Alternative molecular formats and therapeutic applications for bispecific antibodies. Mol Immunol. 2015;67(2):95–106. doi:10.1016/j.molimm.2015.01.003

11. Dumontet C, Reichert JM, Senter PD, Lambert JM, Beck A. Antibody–drug conjugates come of age in oncology. Nat Rev Drug Discov. 2023;22(8):641–661. doi:10.1038/s41573-023-00709-2

12. Joubbi S, Micheli A, Milazzo P, et al. Antibody design using deep learning: from sequence and structure design to affinity maturation. Brief Bioinform. 2024;25(4). doi:10.1093/bib/bbae307

13. Mieczkowski C, Zhang X, Lee D, et al. Blueprint for antibody biologics developability. MAbs. 2023;15(1). doi:10.1080/19420862.2023.2185924

14. Navarro S, Ventura S. Computational methods to predict protein aggregation. Curr Opin Struct Biol. 2022;73:102343. doi:10.1016/j.sbi.2022.102343

15. Fernández-Quintero ML, Ljungars A, Waibl F, et al. Assessing developability early in the discovery process for novel biologics. MAbs. 2023;15(1). doi:10.1080/19420862.2023.2171248

16. Udugama IA, Lopez PC, Gargalo CL, Li X, Bayer C, Gernaey K V. Digital Twin in biomanufacturing: challenges and opportunities towards its implementation. Syst Microbiol Biomanufacturing. 2021;1(3):257–274. doi:10.1007/s43393-021-00024-0

17. Barata TS, Zhang C, Dalby PA, Brocchini S, Zloh M. Identification of protein-excipient interaction hotspots using computational approaches. Int J Mol Sci. 2016;17(6). doi:10.3390/ijms17060853

18. Li J, Wang H, Wang L, Yu D, Zhang X. Stabilization effects of saccharides in protein formulations: A review of sucrose, trehalose, cyclodextrins and dextrans. Eur J Pharm Sci. 2024;192(August 2023):106625. doi:10.1016/j.ejps.2023.106625

19. Sudrik CM, Cloutier T, Mody N, Sathish HA, Trout BL. Understanding the role of preferential exclusion of sugars and polyols from native state IgG1 monoclonal antibodies and its effect on aggregation and reversible self-association. Pharm Res. 2019;36(8):1–12. doi:10.1007/s11095-019-2642-3

20. Cloutier T, Sudrik C, Mody N, Sathish HA, Trout BL. Molecular computations of preferential interaction coefficients of IgG1 monoclonal antibodies with sorbitol, sucrose, and trehalose and the impact of these excipients on aggregation and viscosity. Mol Pharm. 2019;16(8):3657–3664. doi:10.1021/acs.molpharmaceut.9b00545

21. Cloutier TK, Sudrik C, Mody N, Hasige SA, Trout BL. Molecular computations of preferential interactions of proline, arginine.HCl, and NaCl with IgG1 antibodies and their impact on aggregation and viscosity. MAbs. 2020;12(1):1–12. doi:10.1080/19420862.2020.1816312

22. Wang Y, Williams HD, Dikicioglu D, Dalby PA. Predictive Model Building for Aggregation Kinetics Based on Molecular Dynamics Simulations of an Antibody Fragment. Mol Pharm. Published online 2024. doi:10.1021/acs.molpharmaceut.4c00859

23. Rospiccio M, Arsiccio A, Winter G, Pisano R. The role of cyclodextrins against interface-induced denaturation in pharmaceutical formulations: A molecular dynamics approach. Mol Pharm. 2021;18(6):2322–2333. doi:10.1021/acs.molpharmaceut.1c00135

24. Kalayan J, Curtis RA, Warwicker J, Henchman RH. Thermodynamic origin of differential excipient-lysozyme interactions. Front Mol Biosci. 2021;8(June):1–13. doi:10.3389/fmolb.2021.689400

25. Calero-Rubio C, Ghosh R, Saluja A, Roberts CJ. Predicting protein-protein interactions of concentrated antibody solutions using dilute solution data and coarse-grained molecular models. J Pharm Sci. 2018;107(5):1269–1281. doi:10.1016/j.xphs.2017.12.015

26. Wang J, Olsson S, Wehmeyer C, et al. Machine Learning of Coarse-Grained Molecular Dynamics Force Fields. ACS Cent Sci. 2019;5(5):755–767. doi:10.1021/acscentsci.8b00913

27. Cloutier TK, Sudrik C, Mody N, Sathish HA, Trout BL. Machine learning models of antibody-excipient preferential interactions for use in computational formulation design. Mol Pharm. 2020;17(9):3589–3599. doi:10.1021/acs.molpharmaceut.0c00629

28. Jo S, Xu A, Curtis JE, Somani S, Mackerell AD. Characterization of antibody-excipient interactions for rational excipient selection using the site identification by ligand competitive saturation-biologics approach. Mol Pharm. 2020;17(11):4323–4333. doi:10.1021/acs.molpharmaceut.0c00775

29. Zhang C, Gossert ST, Williams J, et al. Ranking mAb–excipient interactions in biologics formulations by NMR spectroscopy and computational approaches. MAbs. 2023;15(1). doi:10.1080/19420862.2023.2212416

30. Li X, Orr AA, Sajadi MM, et al. Investigating the interaction between excipients and monoclonal antibodies PGT121 and N49P9.6-FR-LS: A comprehensive analysis. ChemRxiv. doi:10.26434/chemrxiv-2024-1zv1q

31. Somani S, Jo S, Thirumangalathu R, et al. Toward biotherapeutics formulation composition engineering using site-identification by ligand competitive saturation (SILCS). J Pharm Sci. 2021;110(3):1103–1110. doi:10.1016/j.xphs.2020.10.051

32. Zalar M, Svilenov HL, Golovanov AP. Binding of excipients is a poor predictor for aggregation kinetics of biopharmaceutical proteins. Eur J Pharm Biopharm. 2020;151(December 2019):127–136. doi:10.1016/j.ejpb.2020.04.002

33. Dong J, Wu Z, Xu H, Ouyang D. FormulationAI : A novel web-based platform for drug. Brief Bioinform. 2024;25(1):1–10. 10.1093/bib/bbad419

34. Patel S, Patel M, Kulkarni M, Patel MS. DE-INTERACT: A machine-learning-based predictive tool for the drug-excipient interaction study during product development—Validation through paracetamol and vanillin as a case study. Int J Pharm. 2023;637(March):122839. doi:10.1016/j.ijpharm.2023.122839

35. Rao VA, Kim JJ, Patel DS, Rains K, Estoll CR. A comprehensive scientific survey of excipients used in currently marketed, therapeutic biological drug products. Pharm Res. 2020;37(10). doi:10.1007/s11095-020-02919-4

36. Strickley RG, Lambert WJ. A review of formulations of commercially available antibodies. J Pharm Sci. 2021;110(7):2590-2608.e56. doi:10.1016/j.xphs.2021.03.017

37. Wang SS, Yan Y, Ho K. US FDA-approved therapeutic antibodies with high-concentration formulation: summaries and perspectives. Antib Ther. 2021;4(4):262–273. doi:10.1093/abt/tbab027

38. Jumper J, Evans R, Pritzel A, et al. Highly accurate protein structure prediction with AlphaFold. Nature. 2021;596(7873):583–589. doi:10.1038/s41586-021-03819-2

39. Heinzinger M, Elnaggar A, Wang Y, et al. Modeling aspects of the language of life through transfer-learning protein sequences. BMC Bioinformatics. 2019;20(1):1–17. doi:10.1186/s12859-019-3220-8

40. Clifford JN, Høie MH, Deleuran S, Peters B, Nielsen M, Marcatili P. BepiPred-3.0: Improved B-cell epitope prediction using protein language models. Protein Sci. 2022;31(12):e4497. doi:10.1002/pro.4497

41. Rives A, Meier J, Sercu T, et al. Biological structure and function emerge from scaling unsupervised learning to 250 million protein sequences. Proc Natl Acad Sci U S A. 2021;118(15). doi:10.1073/pnas.2016239118

42. Elnaggar A, Heinzinger M, Dallago C, et al. ProtTrans: Toward understanding the language of life through self-supervised learning. IEEE Trans Pattern Anal Mach Intell. 2022;44(10):7112–7127. doi:10.1109/TPAMI.2021.3095381

43. Littmann M, Heinzinger M, Dallago C, Olenyi T, Rost B. Embeddings from deep learning transfer GO annotations beyond homology. Sci Rep. 2021;11(1):1–14. doi:10.1038/s41598-020-80786-0

44. Cock PJA, Antao T, Chang JT, et al. Biopython: Freely available Python tools for computational molecular biology and bioinformatics. Bioinformatics. 2009;25(11):1422–1423. doi:10.1093/bioinformatics/btp163

45. Geurts P, Ernst D, Wehenkel L. Extremely randomized trees. Mach Learn. 2006;63(1):3–42. doi:10.1007/s10994-006-6226-1

46. Pedregosa F, Varoquaux G, Gramfort A, et al. Scikit-learn: Machine learning in python. J Mach Learn Res. 2011;12:2825–2830. http://scikit-learn.sourceforge.net.

47. Waight AB, Prihoda D, Shrestha R, et al. A machine learning strategy for the identification of key in silico descriptors and prediction models for IgG monoclonal antibody developability properties. MAbs. 2023;15(1). doi:10.1080/19420862.2023.2248671

48. Chawla N V, Bowyer KW, Hall LO, Kegelmeyer WP. SMOTE: synthetic minority over-sampling technique. J Artif Int Res. 2002;16(1):321–357. doi:10.1613/jair.953

49. Fu L, Niu B, Zhu Z, Wu S, Li W. CD-HIT: Accelerated for clustering the next-generation sequencing data. Bioinformatics. 2012;28(23):3150–3152. doi:10.1093/bioinformatics/bts565

50. Li W, Godzik A. CD-HIT: A fast program for clustering and comparing large sets of protein or nucleotide sequences. Bioinformatics. 2006;22(13):1658–1659. doi:10.1093/bioinformatics/btl158

51. Lundberg SM, Lee SI. A unified approach to interpreting model predictions. In: Guyon I, Luxburg U V, Bengio S, et al., eds. Advances in Neural Information Processing Systems 30. Curran Associates, Inc.; 2017:4765–4774. http://papers.nips.cc/paper/7062-a-unified-approach-to-interpreting-model-predictions.pdf

52. Breiman L. Random forests. Mach Learn. 2001;45(1):5–32. doi:10.1023/A:1010933404324

