## Supplementary Figures for "Machine learning driven acceleration of biopharmaceutical formulation development using Excipient Prediction Software (ExPreSo)"

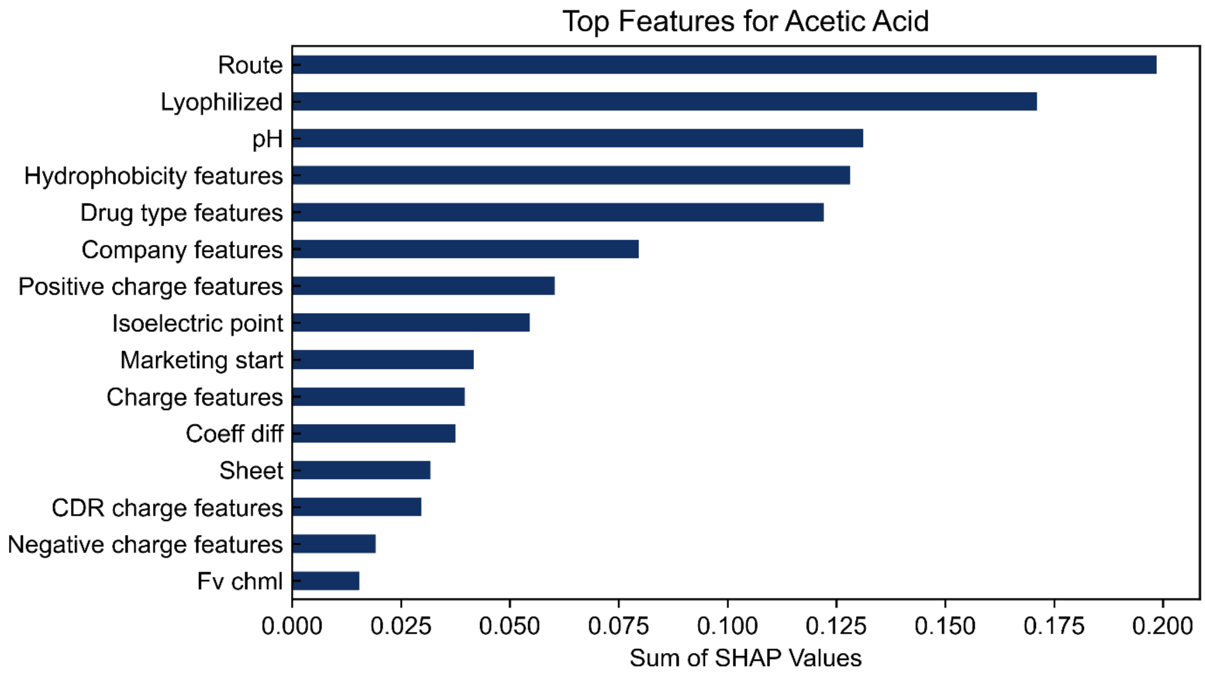


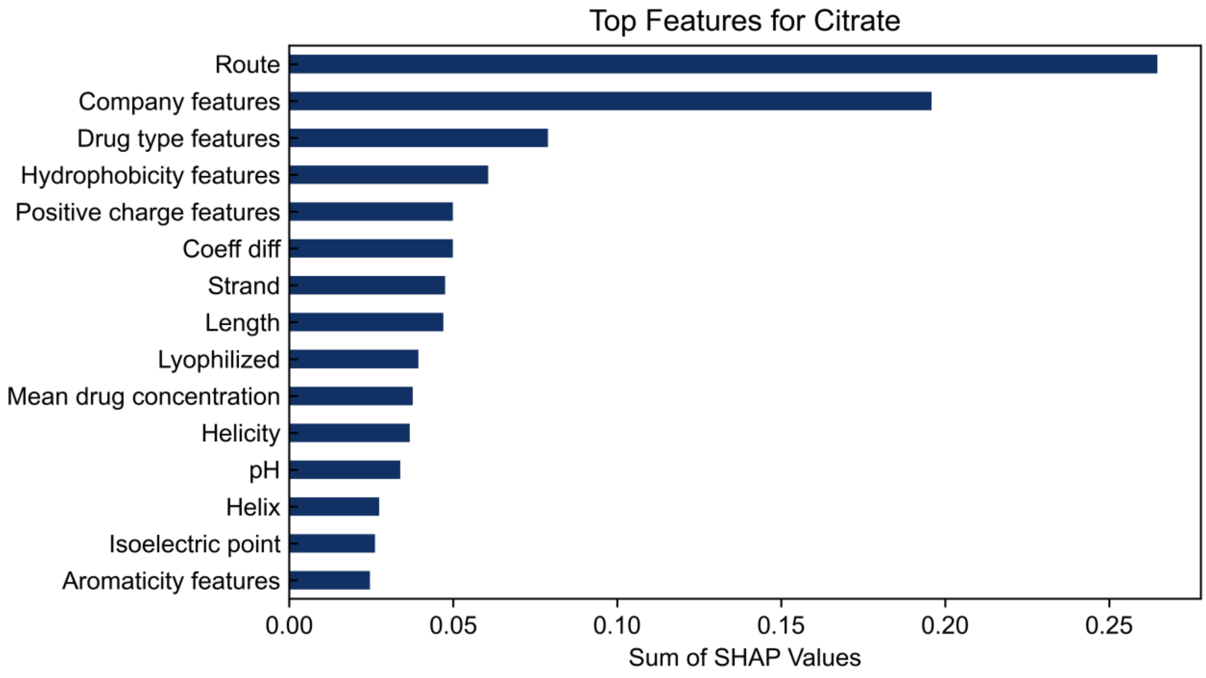

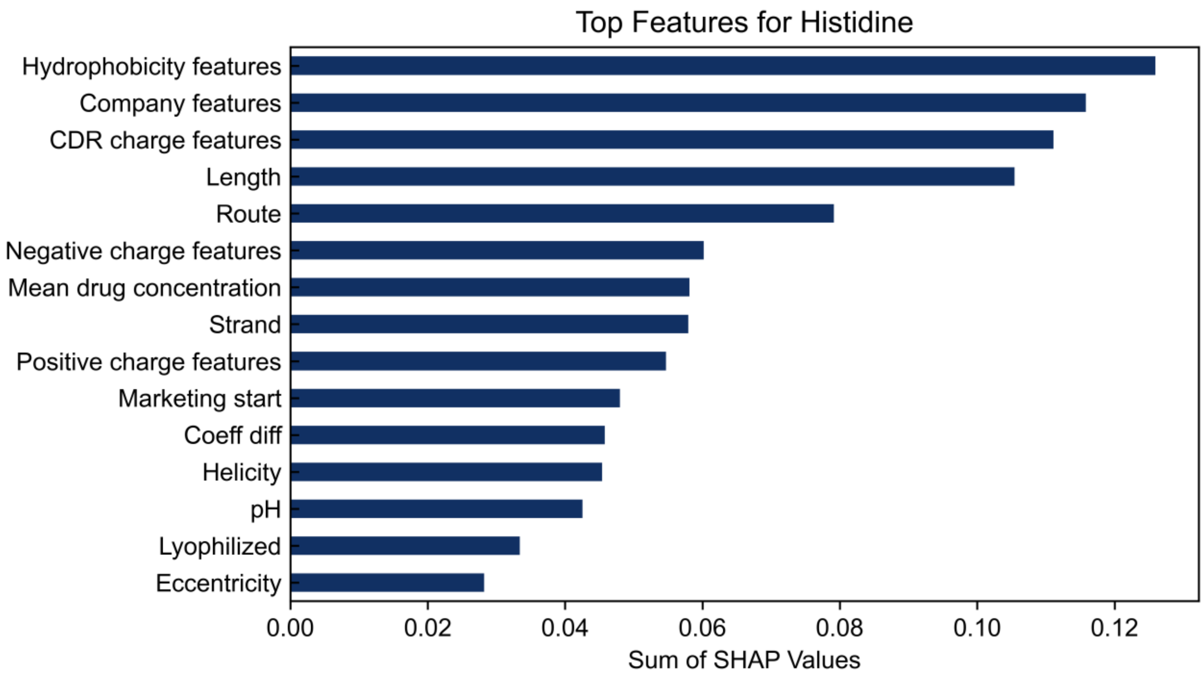

Top Features for Histidine
Hydrophobicity features
Company features
CDR charge features
Length
Route
Negative charge features
Mean drug concentration
Strand
Positive charge features
Marketing start
Coeff diff
Helicity
pH
Lyophilized
Eccentricity
0.00
0.02
0.04
0.06
0.08
0.10
0.12
Sum of SHAP Values


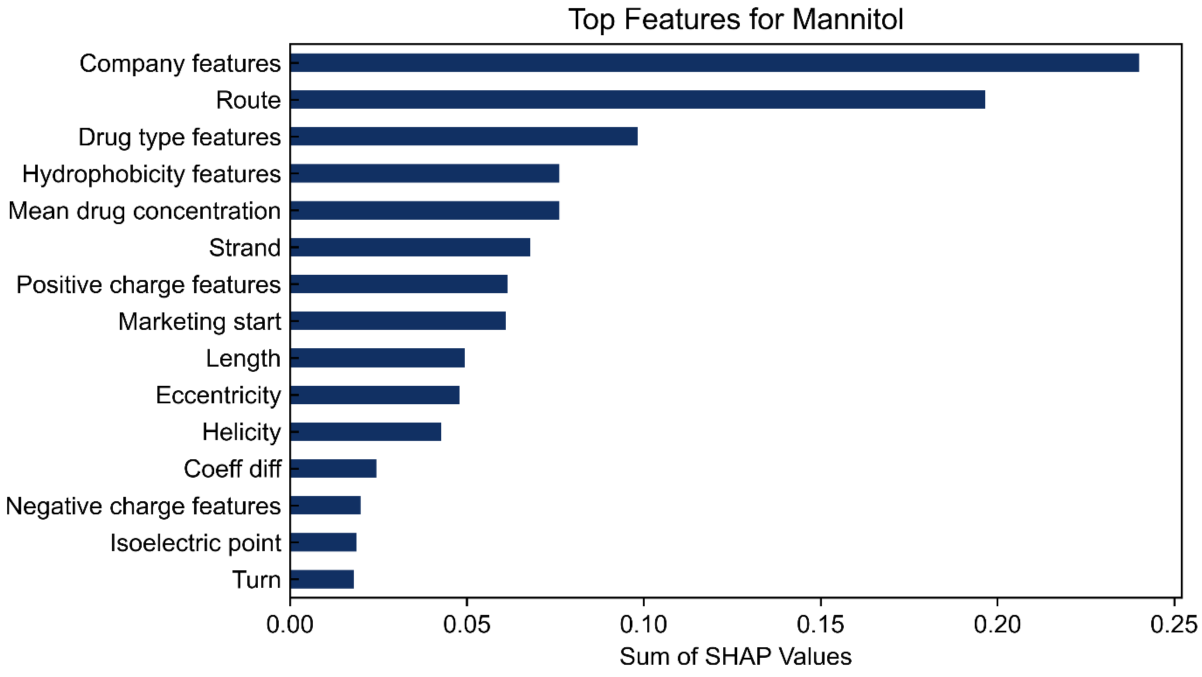

Top Features for Mannitol
Company features
Route
Drug type features
Hydrophobicity features
Mean drug concentration
Strand
Positive charge features
Marketing start
Length
Eccentricity
Helicity
Coeff diff
Negative charge features
Isoelectric point
Turn
0.00
0.05
0.10
0.15
0.20
0.25
Sum of SHAP Values

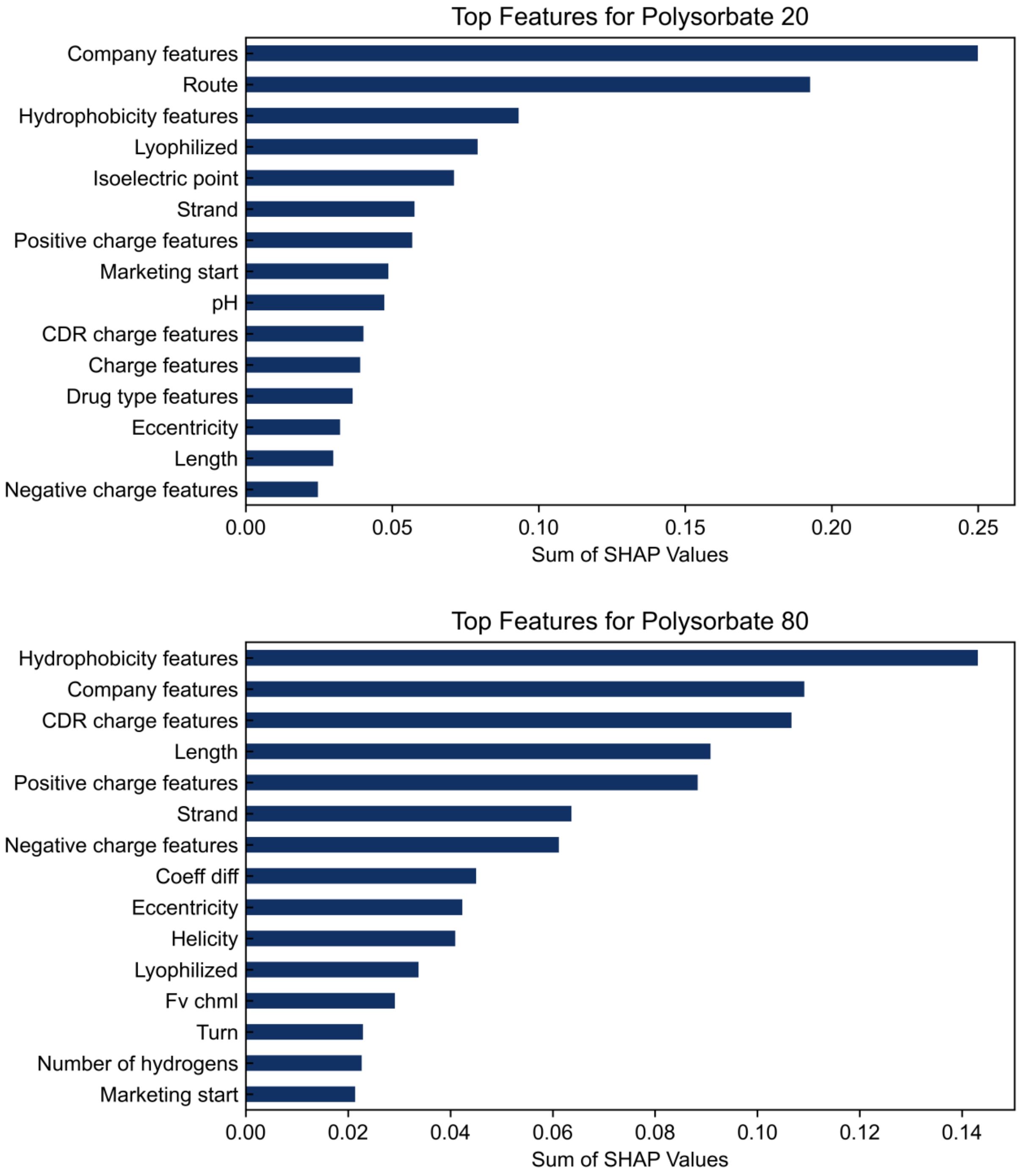

Top Features for Polysorbate 20
Company features
Route
Hydrophobicity features
Lyophilized
Isoelectric point
Strand
Positive charge features
Marketing start
pH
CDR charge features
Charge features
Drug type features
Eccentricity
Length
Negative charge features
0.00
0.05
0.10
0.15
0.20
0.25
Sum of SHAP Values
Top Features for Polysorbate 80
Hydrophobicity features
Company features
CDR charge features
Length
Positive charge features
Strand
Negative charge features
Coeff diff
Eccentricity
Helicity
Lyophilized
Fv chml
Turn
Number of hydrogens
Marketing start
0.00
0.02
0.04
0.06
0.08
0.10
0.12
0.14
Sum of SHAP Values

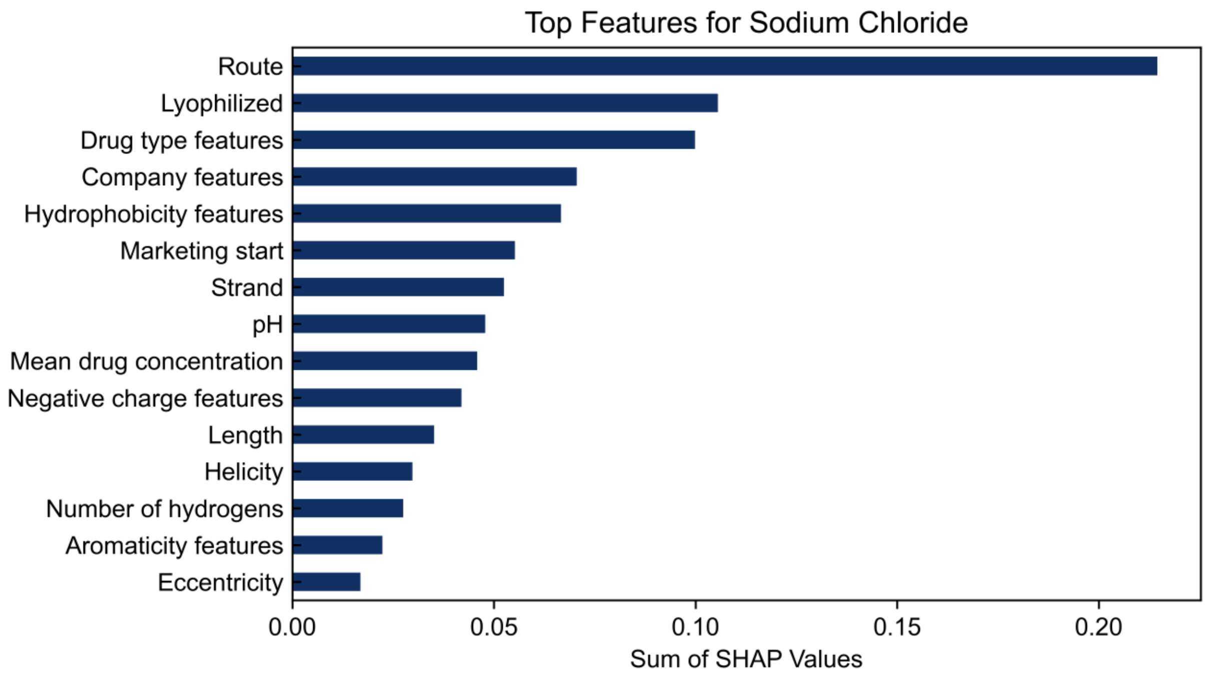

Top Features for Sodium Chloride
Route
Lyophilized
Drug type features
Company features
Hydrophobicity features
Marketing start
Strand
pH
Mean drug concentration
Negative charge features
Length
Helicity
Number of hydrogens
Aromaticity features
Eccentricity
0.00
0.05
0.10
0.15
0.20
Sum of SHAP Values


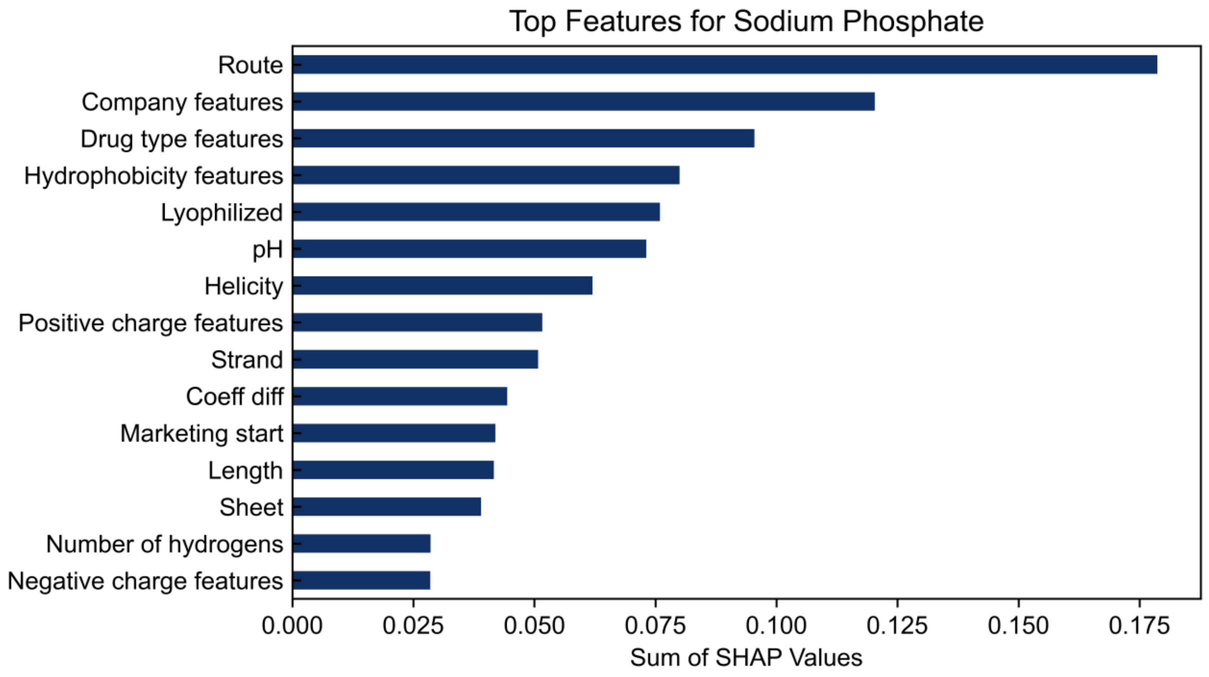

Top Features for Sodium Phosphate
Route
Company features
Drug type features
Hydrophobicity features
Lyophilized
pH
Helicity
Positive charge features
Strand
Coeff diff
Marketing start
Length
Sheet
Number of hydrogens
Negative charge features
0.000
0.025
0.050
0.075
0.100
0.125
0.150
0.175
Sum of SHAP Values

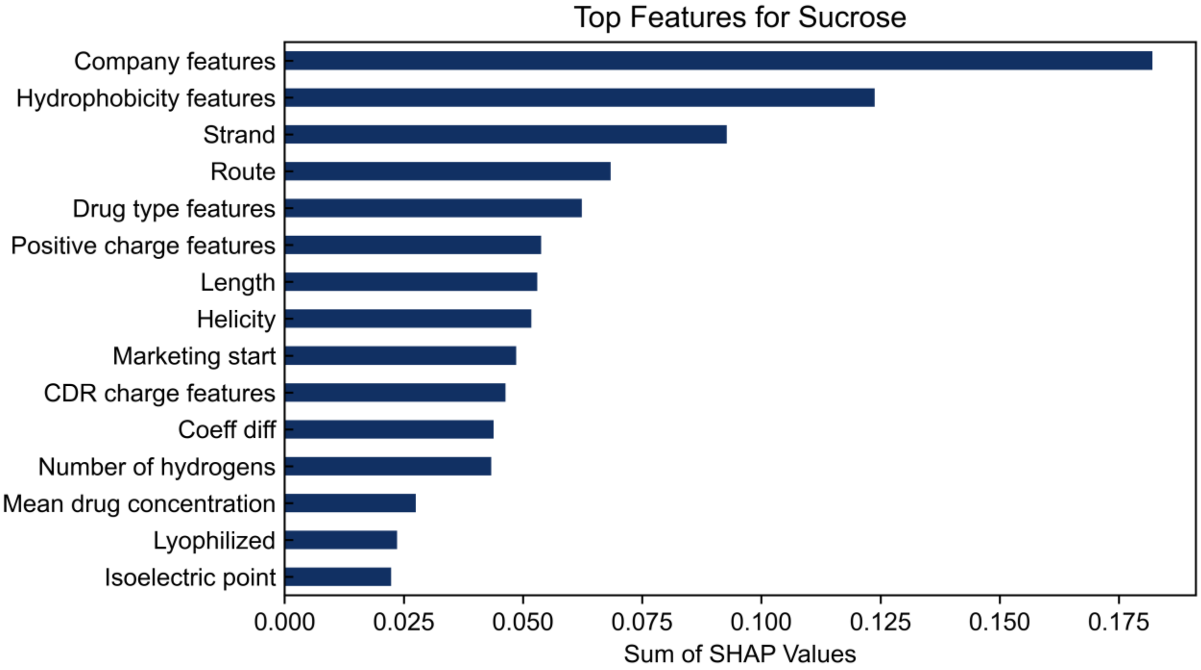


Fig S1: Feature importances within the machine-learning model for the prediction of each excipient in the “Interpretable” version of ExPreSo, which lacks non-interpretable features such as embeddings and amino acid frequencies. The summed absolute SHAP values are shown for each category. Higher values indicate a higher importance in the model. To show broader trends, we grouped most individual features into larger categories, for example the Route category contained the individual boolean features “route:INTRAVENOUS” and “route:SUBCUTANEOUS”.

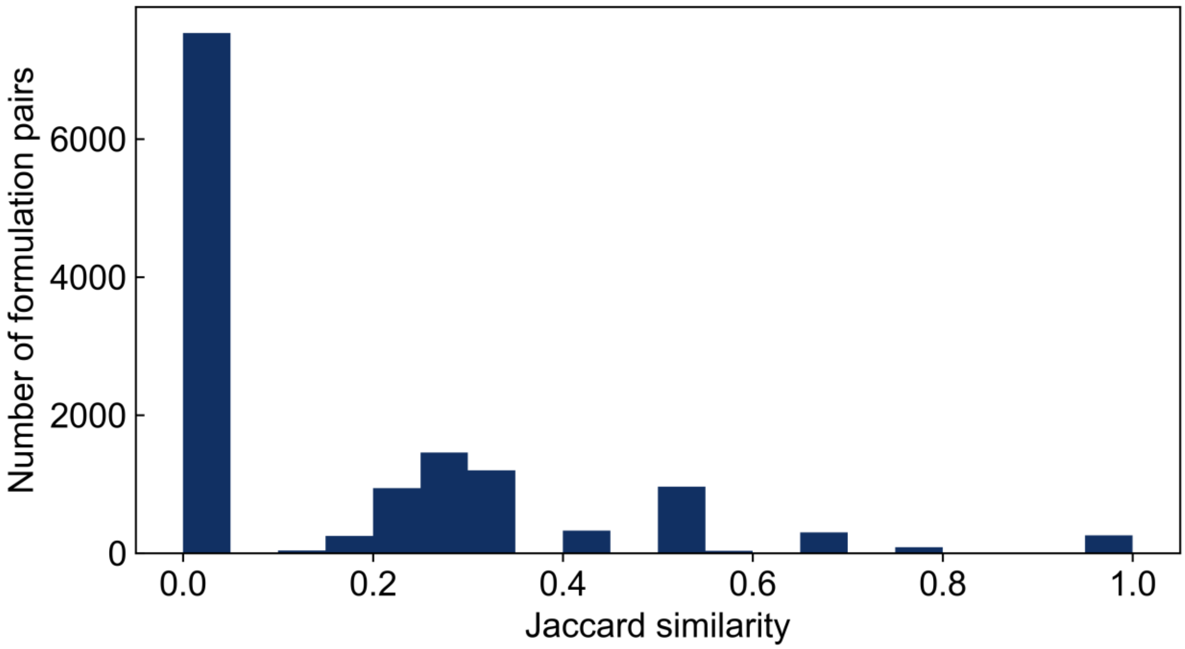


Fig S2: Pairwise similarity of approved therapeutic protein formulations in the ExPreSo dataset as measured with the Jaccard index. A formulation pair with identical excipients has a Jaccard index of 1.0. A formulation pair with no shared excipients has a Jaccard index of 0.0. The Jaccard index was measured for all pairs of formulations that were not in the same cluster used for leave-one-group-out cross-validation (i.e. not closely related sequences).

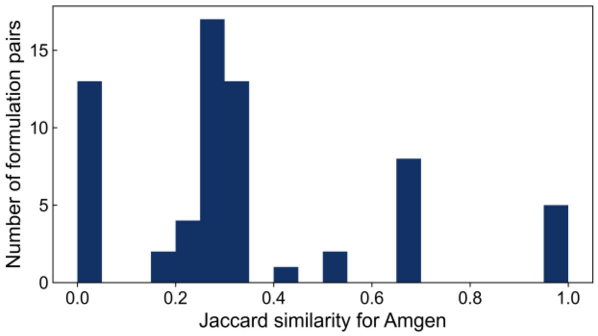
Number of formulation pairs
0
5
10
15
0.0
0.2
0.4
0.6
0.8
1.0
Jaccard similarity for Amgen

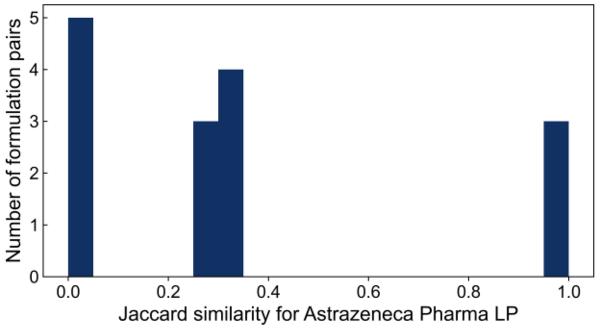
Number of formulation pairs
0
1
2
3
4
5
0.0
0.2
0.4
0.6
0.8
1.0
Jaccard similarity for Astrazeneca Pharma LP

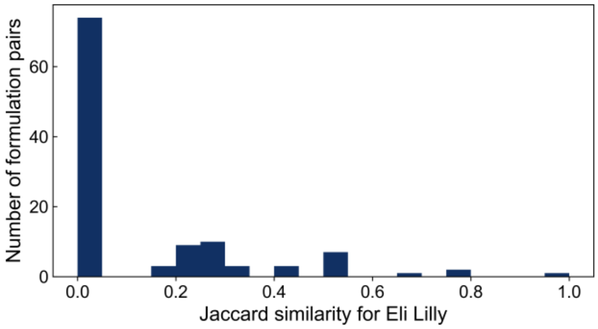
Number of formulation pairs
0
20
40
60
0.0
0.2
0.4
0.6
0.8
1.0
Jaccard similarity for Eli Lilly

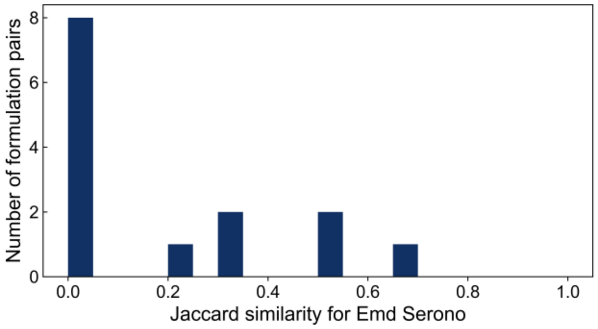
Number of formulation pairs
0
2
4
6
8
0.0
0.2
0.4
0.6
0.8
1.0
Jaccard similarity for Emd Serono

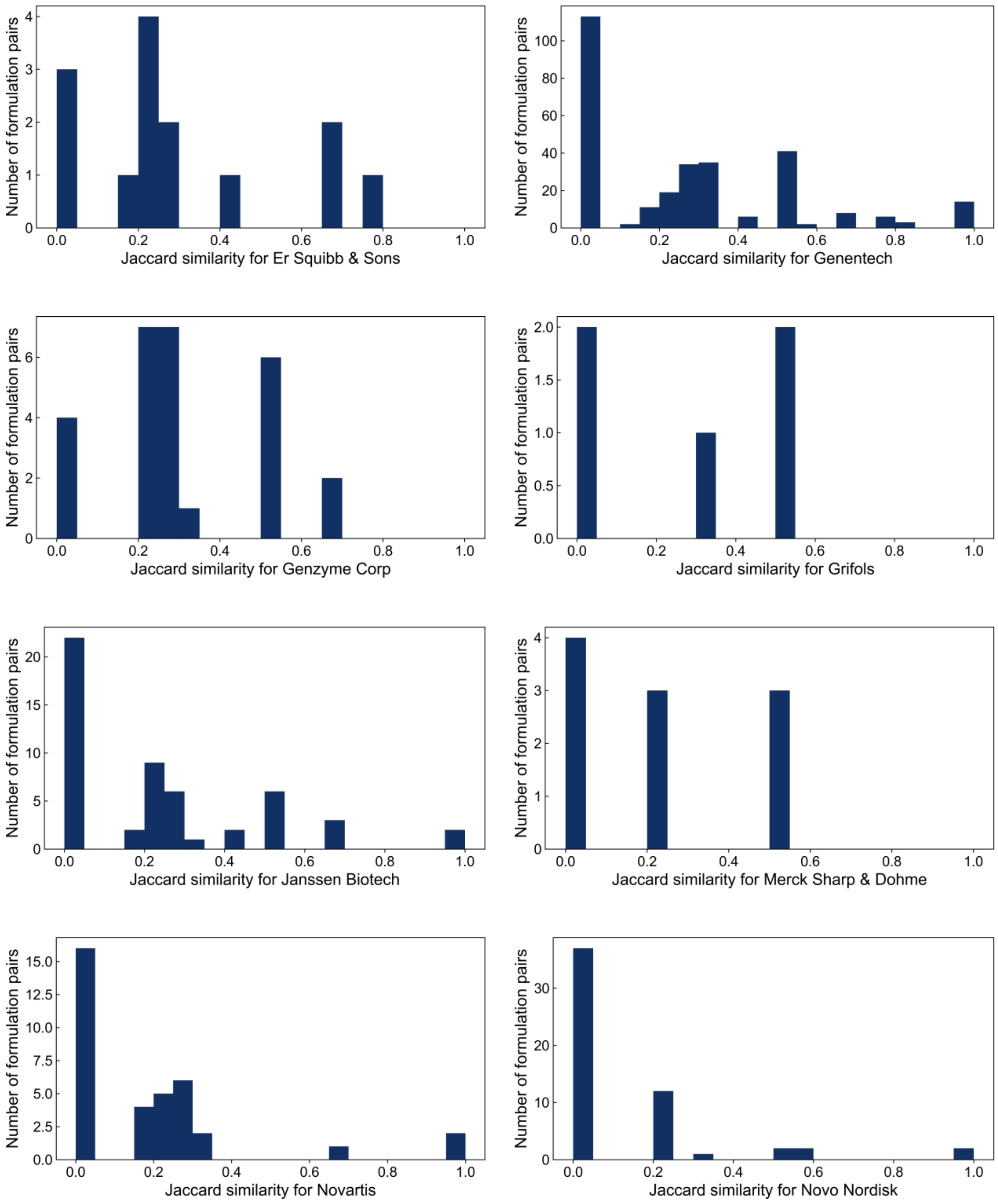
Number of formulation pairs
Jaccard similarity for Er Squibb & Sons
Jaccard similarity for Genentech
Jaccard similarity for Genzyme Corp
Jaccard similarity for Grifols
Jaccard similarity for Janssen Biotech
Jaccard similarity for Merck Sharp & Dohme
Jaccard similarity for Novartis
Jaccard similarity for Novo Nordisk

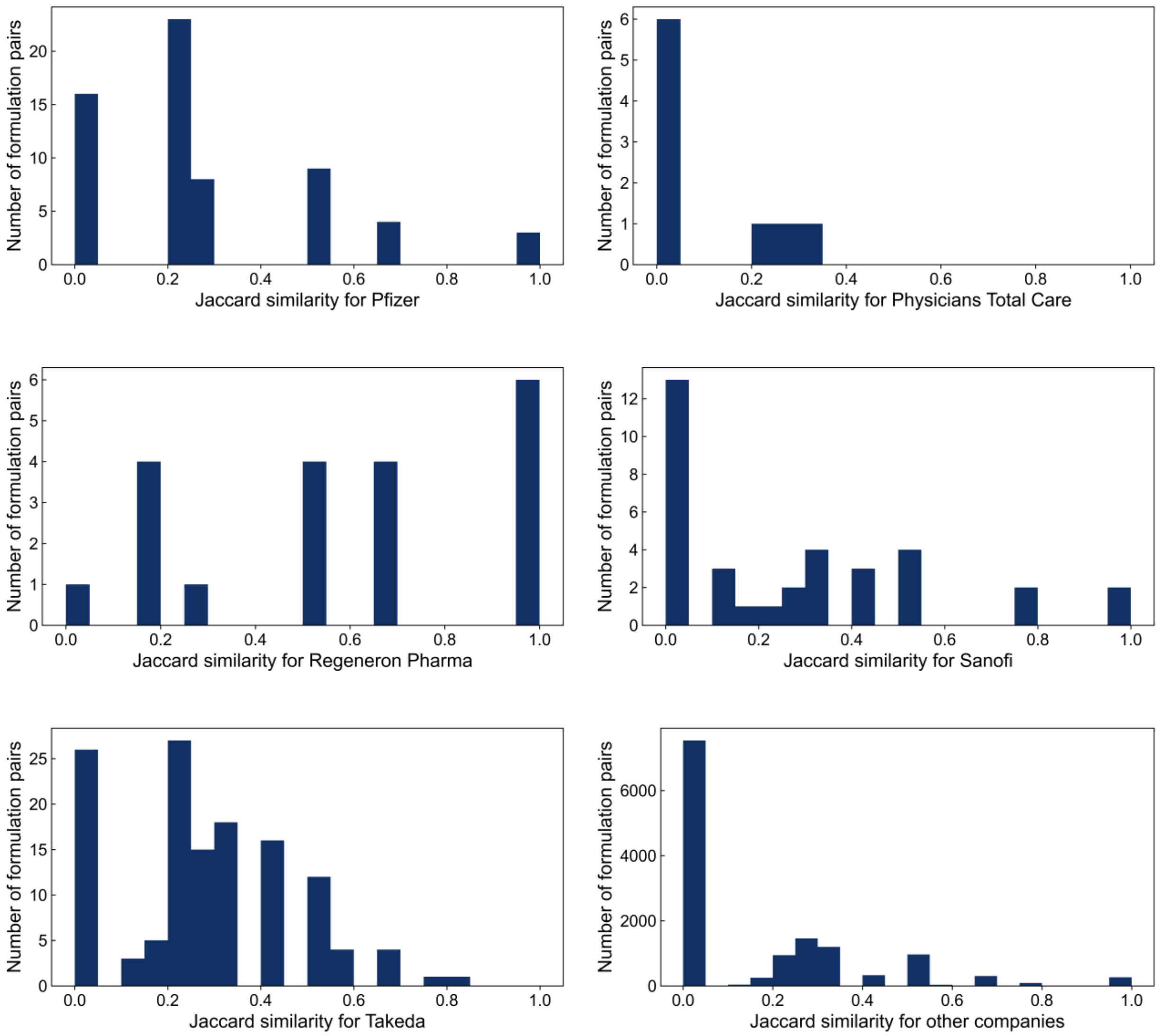


Fig S3. Pairwise similarity of approved therapeutic protein formulations by the top companies in the dataset. A formulation pair with identical excipients has a Jaccard index of 1.0. A formulation pair with no shared excipients has a Jaccard index of 0.0. The Jaccard index was measured for all pairs of formulations that were not in the same cluster used for leave-one-group-out cross-validation (i.e. not closely related sequences).

Table S1. Initial list of features used in ExPreSo

| |
|---|
| 1024 ProtTrans pLM Embeddings |
| 400 Dipeptide frequencies |
| 20 Amino acid frequencies |
| Company: Amgen |
| Company: Astrazeneca Pharma LP |
| Company: Eli Lilly |
| Company: EMD Serono |

| |
|---|
| Company: ER Squibb & Sons |
| Company: Genentech |
| Company: Genzyme Corp |
| Company: Grifols |
| Company: Janssen Biotech |
| Company: Merck Sharp & Dohme |
| Company: Novartis |
| Company: Novo Nordisk |
| Company: Pfizer |
| Company: Physicians Total Care |
| Company: Regeneron Pharma |
| Company: Sanofi |
| Company: Takeda |
| Company: Other Company |
| Route: Intradermal |
| Route: Intramuscular |
| Route: Intravenous |
| Route: Intravitreal |
| Route: Nasal |
| Route: Ophthalmic |
| Route: Oral |
| Route: Parenteral |
| Route: Respiratory (Inhalation) |
| Route: Soft Tissue |
| Route: Subcutaneous |
| Route: Topical |
| Type: IgG1 |
| Type: IgG4 |
| Type: IgG2 |
| Type: mAb |
| Energy |
| Number of Hydrogens |
| Total Hydrophobic Patch Size |
| Largest Hydrophobic Patch |
| Total Area of the 2 largest Hydrophobic Patches |
| Total Area of the 3 largest Hydrophobic Patches |
| Total Area of the 4 largest Hydrophobic Patches |
| Total Area of the 5 largest Hydrophobic Patches |
| Number of Hydrophobic Patches |
| Percentage of Hydrophobic Patches to Total Surface Area |
| Total Positively Charged Patch Size |
| Largest Positively Charged Patch |
| Total Area of the 2 largest Positively Patches |
| Total Area of the 3 largest Positively Patches |
| Total Area of the 4 largest Positively Patches |
| Total Area of the 5 largest Positively Patches |
| Number of Positively Charged Patches |
| Percentage of Positively Charged Patches to Total Surface Area |
| Total Negatively Charged Patch Size |
| Largest Negatively Charged Patch |

| |
|---|
| Total Area of the 2 largest Negatively Patches |
| Total Area of the 3 largest Negatively Patches |
| Total Area of the 4 largest Negatively Patches |
| Total Area of the 5 largest Negatively Patches |
| Number of Negatively Charged Patches |
| Percentage of Negatively Charged Patches to Total Surface Area |
| Total Ionic Patch Size |
| Largest Ionic Patch |
| Total Area of the 2 largest Ionic Patches |
| Total Area of the 3 largest Ionic Patches |
| Total Area of the 4 largest Ionic Patches |
| Total Area of the 5 largest Ionic Patches |
| Number of Ionic Patches |
| Percentage of Ionic Patches to Total Surface Area |
| Total Hydrophobic Patch Size near CDR |
| Largest Hydrophobic Patch near CDR |
| Total Area of the 2 largest Hydrophobic Patches near CDR |
| Total Area of the 3 largest Hydrophobic Patches near CDR |
| Total Area of the 4 largest Hydrophobic Patches near CDR |
| Total Area of the 5 largest Hydrophobic Patches near CDR |
| Number of Hydrophobic Patches near CDR |
| Total Positively Charged Patch Size near CDR |
| Largest Positively Charged Patch near CDR |
| Total Area of the 2 largest Positively Charged Patches near CDR |
| Total Area of the 3 largest Positively Charged Patches near CDR |
| Total Area of the 4 largest Positively Charged Patches near CDR |
| Total Area of the 5 largest Positively Charged Patches near CDR |
| Number of Positively Charged Patches near CDR |
| Total Negatively Charged Patch Size near CDR |
| Largest Negatively Charged Patch near CDR |
| Total Area of the 2 largest Negatively Charged Patches near CDR |
| Total Area of the 3 largest Negatively Charged Patches near CDR |
| Total Area of the 4 largest Negatively Charged Patches near CDR |
| Total Area of the 5 largest Negatively Charged Patches near CDR |
| Number of Negatively Charged Patches near CDR |
| Ionic patches close to the CDR region |
| Largest ionic patch close to the CDR region |
| Total Area of the 2 largest Ionic Patches near CDR |
| Total Area of the 3 largest Ionic Patches near CDR |
| Total Area of the 4 largest Ionic Patches near CDR |
| Total Area of the 5 largest Ionic Patches near CDR |
| Number of Ionic Patches near CDR |
| Protein Mass |
| Coordinate-based pI |
| Extinction Coefficient |
| Frictional Coefficient |
| Diffusion Coefficient |
| Radius of Gyration |
| Hydrodynamic Radius |
| Sedimentation Constant |

| |
|---|
| Eccentricity |
| VdW Surface Area |
| Hydrophobic Surface Area |
| Hydrophilic Surface Area |
| VdW Volume |
| Mobility |
| Helicity |
| Strand |
| Henry Function |
| Net Charge |
| Apparent Charge |
| Dipole Moment |
| Hydrophobicity Moment |
| Fv Charge Separation (VH - VL difference) |
| Zeta Potential |
| Zeta Dipole Moment |
| Zeta Quadrupole Moment |
| Sequence-based pI (MOE) |
| Antibody CDR Length |
| Sequence-based Isoelectric point (ProtParam) |
| Sequence-based Aromaticity |
| Sequence-based Hydrophobicity |
| Sequence-based Helix |
| Sequence-based Turn |
| Sequence-based Sheet |
| Sequence-based Length |
| Lyophilized |
| Marketing start |
| Mean drug concentration |
| pH |

Table S2: Fraction of Covariance explained by Principal Component Analysis (PCA) during feature reduction

| Group | Ratio |
|---|---|
| CDR Negative charge | 0.991 |
| CDR hydrophobicity | 0.993 |
| CDR positive charge | 0.991 |
| Hydrophobicity | 0.825 |
| Negative charge | 0.978 |
| Positive charge | 0.975 |
| Amino acid frequency | 0.856 |
| Dipeptide frequency | 0.761 |
| Protein embeddings | 0.934 |
